# DASTevolve: Evolving executable protein-design programs with large language models

**DOI:** 10.64898/2026.11.25.769346

**Authors:** Yiyang Zhao, Zijun Gao, Jinghao Wang, Kewen Wang, Jihong Chen, Chunbin Gu, Pheng-Ann Heng, Chang-Yu Hsieh

## Abstract

Protein engineering often requires improving one property while preserving others across multiple regions or molecular states. An aggregate score can reveal a poor design yet offer little guidance about which decision to revise. We introduce DASTevolve, which represents a design task as two linked abstract syntax trees. The functional tree specifies objectives and state relationships, and its links to the structural tree identify where edits are permitted for each objective. A large language model (LLM), as inspired by the recent works on coding agents, proposes program revisions from evaluation feedback, and a sequence optimizer searches within the compiled constraints. In an ***α***-synuclein binder-design ablation with 2,000 candidate evaluations per method, the best feasible DASTevolve candidate achieved an ESMFold binder-chain confidence score (pLDDT) of 63.1, compared with 52.9 for the strongest of three controls, a single-tree controller. In an enzyme-scaffold redesign comparison with one retained trajectory per condition, four of the first ten accepted proposals under region- and objective-specific feedback revised editing rules or functional mappings. Neither control trajectory showed such revisions over the same window. Additional case studies illustrate how the representation accommodates antibody design, peptide selectivity, and ligand-responsive design. Together, these results show how an LLM can use evaluation feedback to revise an executable protein design program while preserving a traceable link from functional objectives to individual sequence edits.

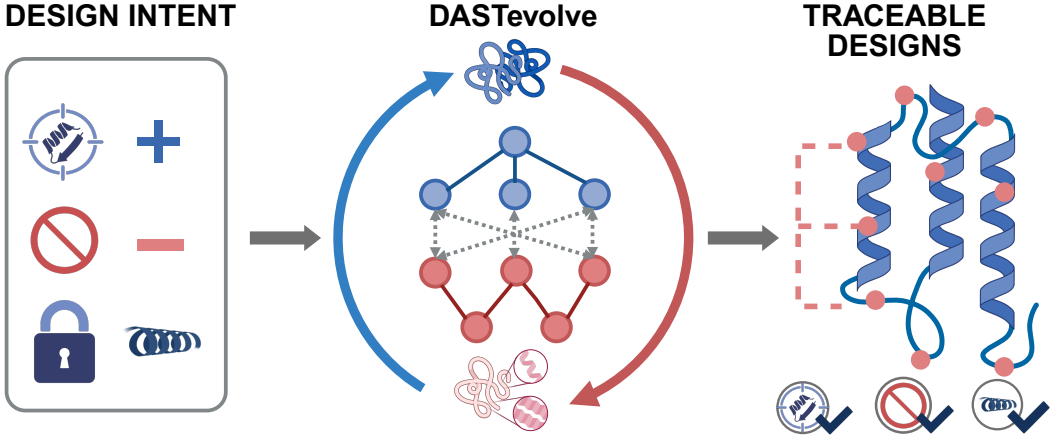

## 1 Introduction

Protein engineering requires translating desired biological functions into a set of testable molecular interventions [1, 2]. A central challenge lies in organizing task-level objectives into coherent intervention hypotheses and continually testing these hypotheses throughout the search process. Advances in sequence generation, structure prediction, and optimization tools provide the computational foundation for executing and evaluating such interventions. Deep learning has improved the evaluation and generation of protein candidates [3–11], expanded the accessible design space, and enabled diverse modes of generation conditioned on sequence, structure, and related information [12–19]. These capabilities are increasingly oriented toward specific application scenarios [20–22] and continue to improve designs through iterative refinement driven by structure-prediction feedback [23, 24].

The knowledge integration, hypothesis generation, and feedback interpretation capabilities of large language models (LLMs) [25] offer a new interface for organizing these decisions. Reasoning-guided and multimodal approaches are beginning to connect natural language with protein sequences, structures, and residue-level design decisions [26, 27]. Recent work has shown that composing and orchestrating existing models and tools under AI-agent scheduling can close the loop of iterative computational protein design workflows [28], and language-guided or multimodal methods have attempted to interpret or generate protein sequences and structures from natural-language intent [29, 30]. However, the ability to invoke tools does not by itself confer well-defined execution semantics on design intent. Sequences and atomic coordinates describe the molecule itself, yet they do not specify what may be changed, what must be preserved, or how molecular states should be compared within a given task, nor does an aggregate candidate score necessarily indicate which intervention hypothesis requires revision [31, 32]. An LLM-facing interface must therefore explicitly connect task objectives, localized operations, and evaluation evidence.

Executable abstraction provides a design principle for establishing this connection: design hypotheses are expressed through task-relevant components and relations, while the permitted transformations, the constraints that must be satisfied, and the evidence required to evaluate outcomes are made explicit. Prior work on programmable protein design has organized complex design requirements into executable programs through modular building blocks, hierarchical constraints, and composable model interfaces [33–35]. For protein engineering, such abstractions must simultaneously express functional objectives and state relations on the one hand, and structural regions and editing permissions on the other. Because a single functional objective may span multiple structural regions and a single region may participate in multiple functions or states, the correspondence between the two must be represented explicitly. Compiling these descriptions into checkable operation scopes, protection conditions, and evaluation rules allows an LLM to propose interventions at the level of task semantics, while specialized molecular tools carry out the underlying computations, preserving a traceable link from design rationale to execution outcome.

An executable representation must also be revisable in light of evidence. A search that falls short of expectations may stem from poor candidate sequences, but it may equally reflect an inappropriate objective decomposition, structural scope, or intervention strategy. AlphaEvolve demonstrated a paradigm in which language models propose program modifications that are then evolved under the guidance of executable evaluators [36], and self-improving agents such as the Darwin Gödel Machine have further explored how the very process of generating successive solutions can improve with feedback [37]. More recently, EnzymeRSI introduced dual-loop improvement at the workflow and algorithm levels for enzyme engineering [38]. For protein design, this line of thinking implies bringing the design representation itself into the optimization loop. The program specifies which interventions may be executed and how their outcomes are evaluated, while evidence from sequence search in turn guides program revision. Such program-level adaptation must be governed by explicit biological constraints, so that design hypotheses and search strategies can adapt while the functional and structural requirements that must be satisfied remain intact.

We present DASTevolve, an RSI-inspired hierarchical protein-program evolution framework organized around three goals (Fig. 1): making protein design mechanism-aware, constraint-preserving, and adaptive. The framework renders the functional decomposition, intervention logic, and supporting evidence of a protein engineering project explicit and revisable, while maintaining biological constraints throughout optimization. We evaluate DASTevolve on fitness optimization, interface design, sequence recovery, redesign under active-site preservation, multistate negative design, lig- and discrimination, and allosteric control. These studies examine whether program revision improves adaptability, preserves biological constraints, and supports failure diagnosis. Taken together, the results provide evidence that optimizing executable design programs alongside protein sequences offers a complementary path toward adaptive and traceable protein engineering.

**Fig. 1.**
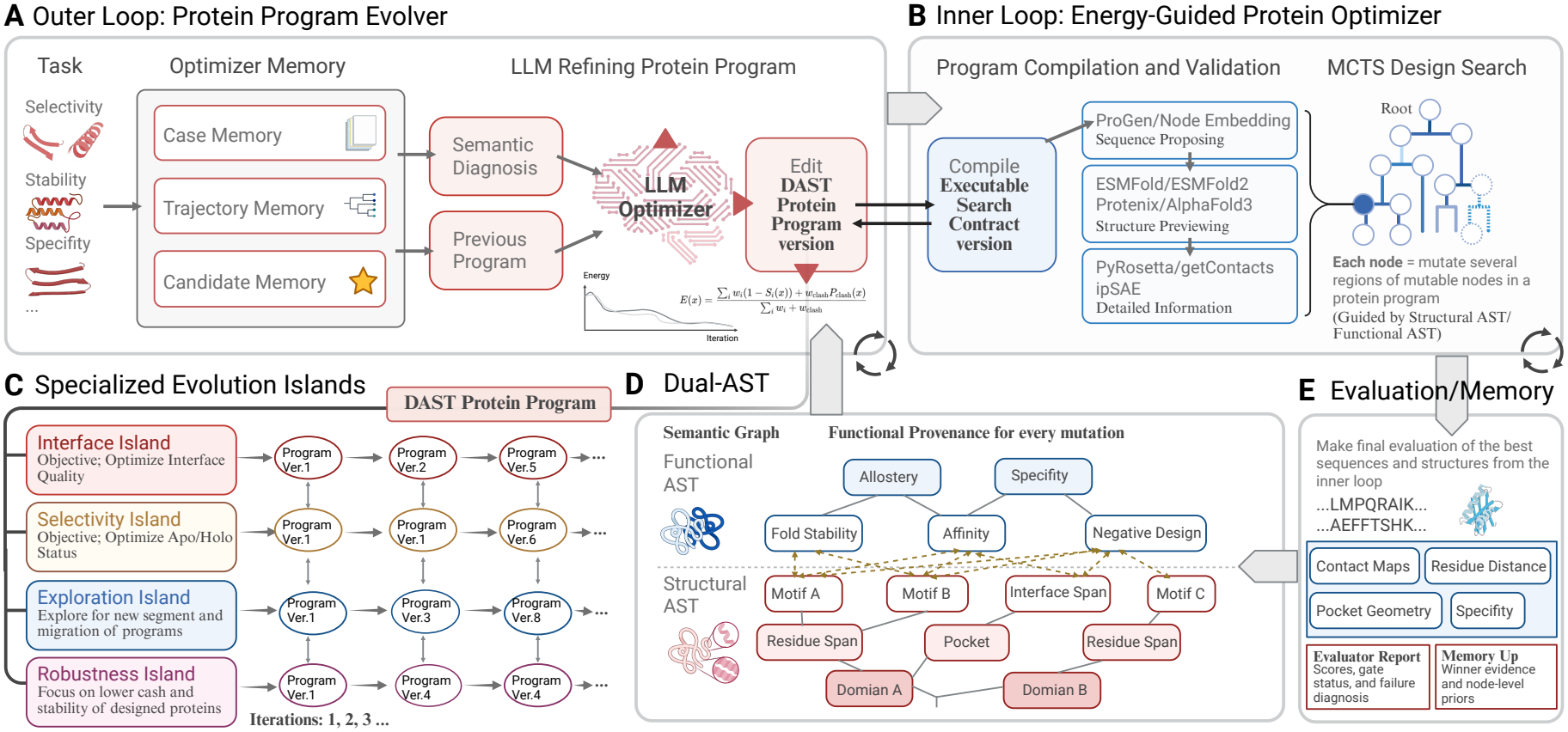
Overview of the DASTevolve framework for hierarchical protein-program evolution. **(A)** The outer loop represents each task as an executable protein program and proposes bounded, evidence-informed revisions. **(B)** Valid revisions are compiled into search contracts that initialize MCTS with interchangeable proposal, structure, and interaction models. **(C)** Specialized program populations can evolve in parallel within islands emphasizing different responsibilities, with controlled migration of promising programs and evidence. **(D)** The DAST connects functional states and objectives to editable structural regions through typed mappings that preserve mutation provenance. **(E)** Candidates returned by the inner search are assessed by a multi-objective evaluator that reports normalized scores, evidence availability, gate outcomes, and failure diagnoses. Eligible winner evidence and node-level priors can subsequently be written to optimizer memory.

## 2 Results

We first describe how the Dual-AST framework translates design objectives into executable sequence-search programs. We then examine three connected questions. First, can program revision improve adaptive search and design performance under a fixed evaluation budget? Second, how can task-specific knowledge and evaluator evidence guide sequence exploration and subsequent program revisions? Third, can the same representation coordinate desired, competing, and preserved molecular states? This progression follows the design program from executable specification through evidence-guided adaptation to multistate design.

### 2.1 An executable framework for hierarchical protein-program evolution

DASTevolve couples two optimization processes: an outer loop that evolves executable design hypotheses and an inner loop that searches for molecular candidates under each hypothesis. The outer loop specifies structural regions, functional objectives, constraints, and permitted interventions. The inner loop optimizes candidate sequences within this specification and returns designs together with structured evaluation evidence. This feedback guides program selection and revision, allowing design specifications and molecular candidates to improve jointly.

Each task begins with a machine-readable case specification defining input chains, biological states, editable and protected regions, design objectives, and evaluator bindings (Fig. 1A). The resulting protein program is represented by a dual abstract syntax tree (Dual-AST; Fig. 1D). The functional AST specifies states and objectives to promote, suppress, or preserve, together with the evidence required to evaluate them. The structural AST defines the domains, motifs, residue spans, pockets, interfaces, and other regions on which interventions may act. Typed cross-tree mappings connect functional intent to structurally localized operations, preserving the functional and structural provenance of proposed edits.

The outer loop integrates the current program, case knowledge, previous search trajectories, candidate evidence, and evaluator-derived failure diagnoses to propose typed revisions to the program’s representation of functional states, structural selectors, intervention permissions, inner-search evaluator configurations, or search strategies. Accepted revisions produce versioned programs with explicit parent–child records. Before sequence optimization, each proposed program is validated and compiled into an executable search contract (Fig. 1B). Compilation resolves structural selectors into residue masks, identifies immutable positions, instantiates permitted operators and budgets, binds functional states to evaluators, and constructs feasibility gates. Revisions that violate the parent program’s edit envelope, protected structural invariants, or schema and type requirements are rejected.

Each valid contract initializes a fresh, budget-limited MCTS tree whose nodes represent sequence candidates and whose edges represent authorized interventions. Pretrained models and structural analysis tools supply proposals and evaluation evidence through standardized interfaces. Protein language models provide sequence scores and proposal priors, while fast predictors support preliminary structural screening, and more computationally demanding predictors provide higher-fidelity structural and multimer evidence. Physics- and geometry-based tools, including PyRosetta, FoldX, and interface analyses, can supply complementary evidence on stability and molecular interactions. The interfaces standardize inputs, outputs, availability, and failure states, allowing backends to be replaced or upgraded without changing case specifications or program semantics. Candidates can consequently be evaluated progressively across cost–fidelity tiers.

Outer-loop population search draws inspiration from the evaluator-guided program evolution exemplified by AlphaEvolve [36] and adapts OpenEvolve’s open-source population and island management mechanisms [39]. Multiple islands maintain programs in the same representation while applying case-configurable selection priorities, such as interface and fold quality, positive–negative state separation, exploration of alternative structural regions, or robustness against unfavorable states (Fig. 1C). All islands share the same validation rules, protected invariants, and contract-level feasibility requirements. At configured intervals, eligible high-performing programs are proposed for migration and assessed against the destination island’s selection criteria. Versioned programs migrate with their evaluation records and provenance, enabling useful revisions to be exchanged while preserving distinct search trajectories.

Inner-loop candidates undergo multiobjective evaluation and explicit feasibility testing (Fig. 1E), producing two complementary outputs. Evaluator reports record normalized scores, evidence availability, gate outcomes, and failure diagnoses for candidate selection and subsequent program revision. Bounded memory proposals contain only eligible evidence associated with the selected winner, including candidate-level observations and node-level priors approved for reuse. This separation keeps incomplete or failed evaluations out of persistent memory and closes the feedback loop from sequence-level outcomes to program evolution.

The resulting constrained bilevel search maintains program revision and sequence intervention as distinct, coupled optimization objects (Fig. 2). Within each inner search, the compiled contract remains fixed. Higher-fidelity checkpoint evidence may update candidate value estimates, while visit counts remain local to the current program’s MCTS tree. Linking candidate outcomes to program versions and edit constraints makes both biological hypotheses and sequence-level tests traceable, supporting subsequent revision and failure analysis.

**Fig. 2.**
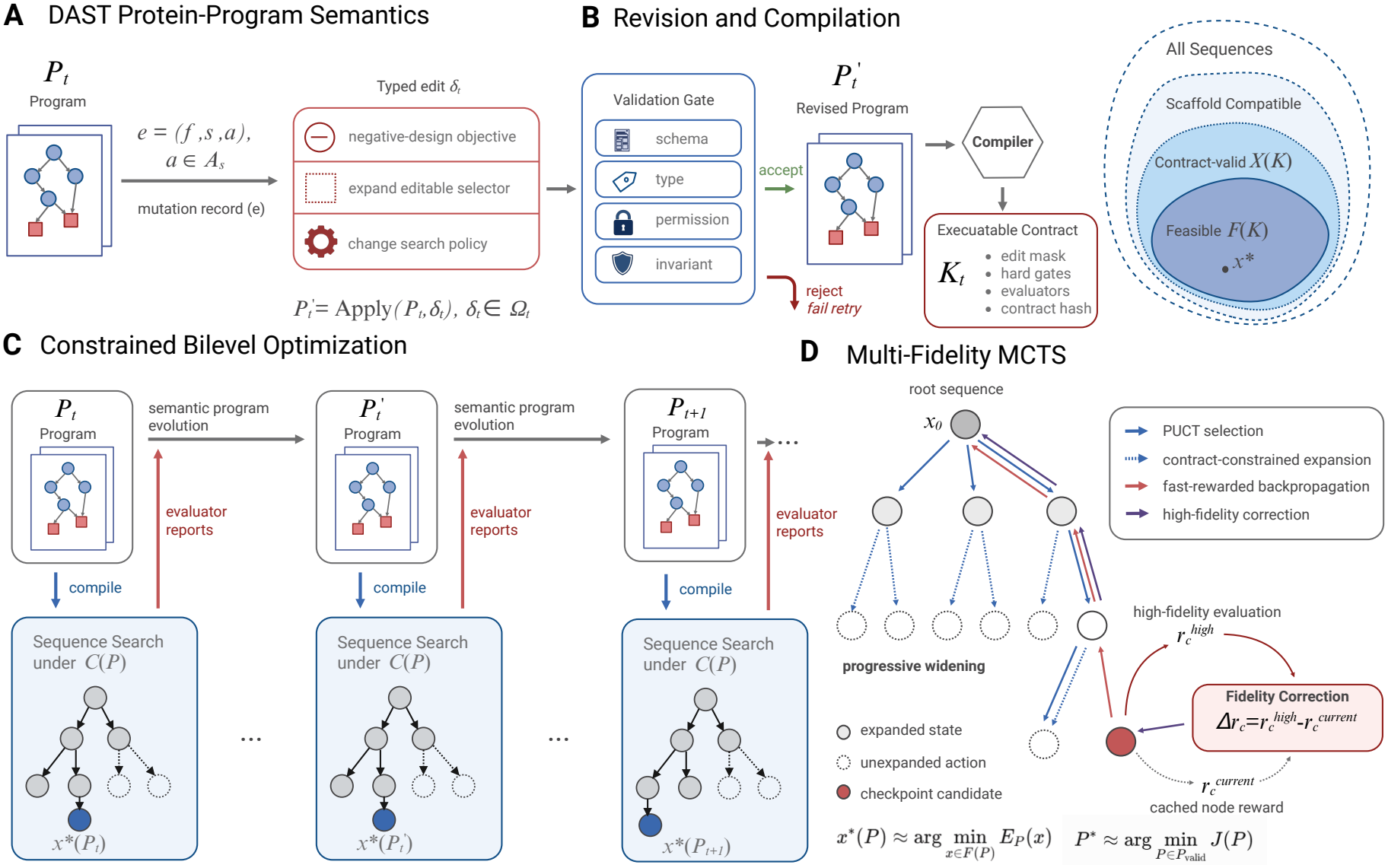
Execution of DASTevolve as constrained bilevel protein-program evolution. **(A)** At outer iteration *t*, the task is represented by a Dual-AST program *P*_*t*_. Typed edges *e* = (*f, s, a*) connect functional requirements to structural scopes and permitted interventions. A bounded edit *δ*_*t*_ *∈*Ω_*t*_ produces a traceable revision 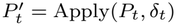. **(B)** Revisions pass schema, type, permission, and invariant checks before being compiled into a search contract 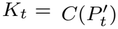. The contract defines editable residues, feasibility gates, evaluator bindings, and provenance; invalid revisions are rejected. It restricts sequence search to the feasible space *ℱ* (*K*_*t*_). **(C)** Each valid program initiates a finite-budget inner search, *x*^*\**^(*P*) *≈*arg min_*x ∈ℱ* (*C*(*P*))_ *E*_*P*_ (*x*), while the outer loop compares programs by the candidates they produce, 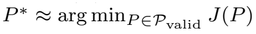 , where *J* (*P*) = *E*_out_(*x*^*\**^(*P*); Ξ) uses the same case-owned evaluation specification Ξ for every competing program within a task and evaluation phase. Evaluator feedback guides subsequent program revisions. **(D)** The inner loop combines PUCT selection, progressive widening, and contract-constrained expansion. Fast rewards support standard MCTS backpropagation, whereas checkpoint candidates receive high-fidelity reevaluation. The correction 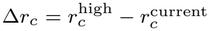 updates accumulated values *W* along the ancestral path without changing visit counts *N*.

### 2.2 Benchmarking adaptive search and matched-budget design performance

We first examine the mechanism of DASTevolve on a large-scale dataset grounded in experimental results from TrpB, and then compare its performance against BAGEL, a programmable protein engineering model of the same type, on the task of designing *α*-synuclein (ASYN)-binding proteins under an identical budget and the same number of iteration rounds. The two experiments respectively evaluate the rationality and effectiveness of the DAST mechanism, and whether DAST-guided structure- and function-oriented protein engineering optimization is more effective than optimization search that compiles energy terms alone. We also conduct an ASYN ablation comparing Full DAST, Static DAST, Single AST, and Flat LLM, to assess the contributions of program adaptation and the dual-AST representation.

We evaluated executable Dual-AST revision in a controlled sequence-selection benchmark derived from experimentally measured TrpB sequence–fitness data [40]. Using the processed by_position split of FLIP2 v3, the archive-based protocol retained 31,557 historical observations as the archive and 7,889 validation sequences as the query pool. In each episode, all strategies used the same sequence sets, optimized the same phase-specific objective, and were limited to 16 queries per round over 12 rounds. We repeated the comparison across four episode seeds. The predefined objective combined experimental fitness with a modular mutation score. The modular component assigned interaction scores to mutation pairs based on their structural and functional node memberships and amino-acid categories, thereby inducing an epistatic landscape. Both phase-specific scoring programs were fixed before sequential selection, ensuring that candidate evaluation was independent of the AST used by each controller.

We compared seven controllers: Full DAST + LLM, Oracle DAST, Flat LLM (flat-feature ridge), Onehot Ridge, Single AST + LLM (one-to-one mapping control), Static DAST, and Random (Fig. 3A). Six used ridge regression to predict the composite objective, while Random sampled unqueried sequences uniformly. Full DAST + LLM started with perturbed structural assignments while leaving the functional mappings intact. In Phase A, patch generation relied on mutation co-occurrence evidence and retained node-identifier semantics. In Phase B, feedback explicitly provided the updated structural assignments and functional mappings. For each phase, we generated four LLM proposals and selected one patch that exactly matched the reference AST, then reused the selected patch across episodes. Full DAST + LLM applied these patches in Rounds 2 and 7. Oracle DAST used the phase-specific reference AST from the outset, whereas Static DAST retained the initial AST. The Single AST + LLM used the patched structural assignments from Round 1, assigned one functional node to each structural node, and switched to the Phase-B assignments in Round 7. Flat LLM and Onehot Ridge relied on AST-independent features.

**Fig. 3.**
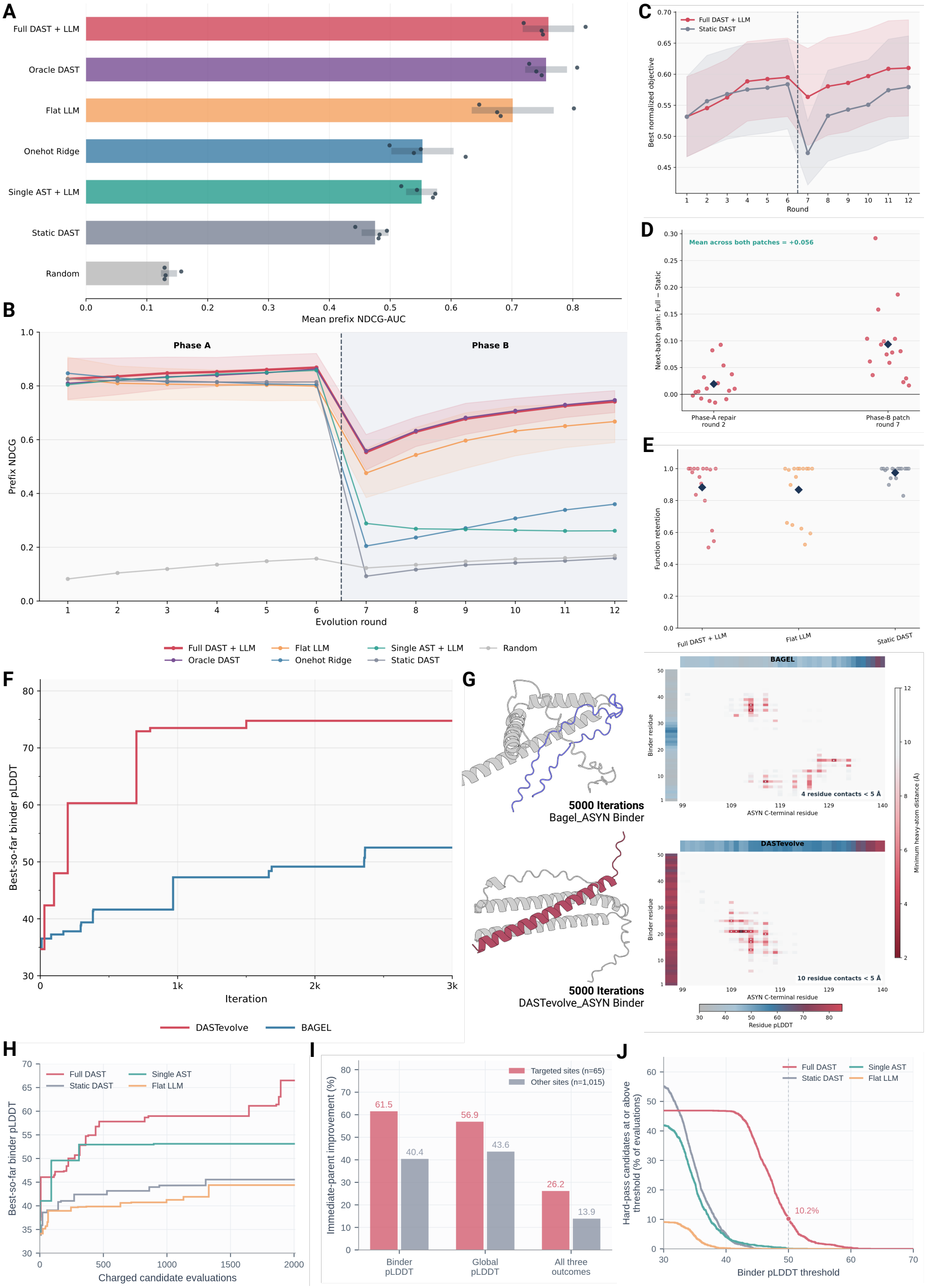
Adaptive TrpB search and ASYN binder design with executable program evolution. **(A)** TrpB prefix NDCG–AUC across selection strategies; points denote seeds and intervals indicate 95% confidence intervals. **(B**,**C)** Ranking quality and best normalized objective across the objective switch between Rounds 6 and 7; C shows the 16-seed mean with 95% confidence intervals. **(D**,**E)** Immediate gains following program repair/patching and retention of the Phase-A function. Points denote seeds and diamonds denote means. **(F**,**G)** Matched-budget ASYN design trajectories (F) and final predicted complexes after 5,000 evaluations (G). Panel F displays the first 3,000 evaluations, as the remaining 2,000 yielded no crucial improvement in best-so-far binder pLDDT. Contact maps show binder–target minimum heavy-atom distances and marginal pLDDT. **(H)** Four-arm ASYN ablation over all 2,000 charged evaluations per arm. **(I)** Immediate-parent improvement frequencies for single substitutions at newly targeted sites (*n* = 65) and other sites (*n* = 1,015) in Full DAST. The joint outcome requires increased binder and global pLDDT and decreased selection loss. **(J)** Fraction of all evaluations passing the complete hard gate and reaching each binder-pLDDT threshold.

At the phase transition, Full DAST + LLM, Oracle DAST, Flat LLM, and Onehot Ridge received base-training labels recomputed under the current objective and used only query history from the current phase. By contrast, Static DAST and the Single AST + LLM retained their Phase-A base labels and accumulated query observations with their originally observed targets across both phases. For all controllers, newly queried candidates were evaluated under the current phase-specific objective. The benchmark therefore assesses reference-assisted adaptation at the level of complete policies that differ in both representation and evidence-update rules. We summarized ranking performance using prefix NDCG–AUC, defined as the mean of the 12 round-wise, within-phase NDCG values. Full DAST + LLM achieved 0.760, compared with 0.756 for Oracle DAST and 0.701 for Flat LLM, and outperformed Flat LLM in all four episodes (Fig. 3A). The remaining scores were 0.553 for Onehot Ridge, 0.552 for the Single AST + LLM, 0.475 for Static DAST, and 0.136 for Random. At the transition before Round 7, the modular assignments and interaction coefficients changed, whereas experimental fitness remained fixed and the within-phase query prefix was reset. Full DAST + LLM and Static DAST ended Phase A with mean prefix NDCG values of 0.868 and 0.814, respectively. During Phase B, Full DAST + LLM improved from 0.553 to 0.742 by Round 12, whereas Static DAST increased only from 0.093 to 0.160 (Fig. 3B). This divergence supports the value of jointly updating the mappings and refreshing the training evidence.

We also evaluated a complementary initial-panel protocol with 16 episode seeds. Each episode began with 32 common observations drawn from the combined 39,446 records, leaving 39,414 candidates available for querying. Objective components were standardized using the initial panel, while controller policies and the 192-query budget remained unchanged. Full DAST + LLM improved sample-efficiency AUC over Static DAST by 0.026, with sample-efficiency AUC defined as the mean of the 12 round-wise best normalized objective values (Fig. 3C). At the revision points in Rounds 2 and 7, Full DAST + LLM achieved a pooled improvement of 0.056 in batch-mean normalized objective over Static DAST across 32 paired episode-by-revision comparisons (Fig. 3D). Full DAST + LLM also achieved a mean baseline-adjusted experimental-fitness retention of 0.883 (Fig. 3E). This metric was defined as the ratio of the best experimental fitness queried in Phase B to that in Phase A after subtracting the pool minimum, with the ratio clipped to [0,1] . Taken together, these results demonstrate the value of combining executable program updates with adaptive candidate selection under controlled changes in modular objectives while continuing to track experimental fitness across the phase transition. This evidence motivated us to further evaluate DASTevolve in ASYN binder design through iterative sequence generation and structure-based evaluation.

We next compared DASTevolve with BAGEL on the same 50-residue ASYN binder-design task [34]. The two methods used a common initial binder, the same fixed target and binder lengths, ESMFold evaluation, and a budget of 5,000 scored designs. This comparison measures computational design performance at a matched molecular-evaluation budget. The target presents a deliberately difficult structural setting: monomeric *α*-synuclein remains intrinsically disordered in mammalian cells, and both structure-prediction and molecular-simulation studies emphasize its heterogeneous conformational ensemble [41, 42]. A single predicted complex therefore cannot represent the full target ensemble. Both methods started from the same randomly generated 50-residue binder, kept the 140-residue ASYN target and binder length fixed, and used ESMFold as the downstream structure predictor [7]. BAGEL performed 5,000 optimization steps, whereas DASTevolve performed 50 outer program-evolution iterations with 100 inner sequence evaluations per iteration, yielding the same total budget of 5,000 scored designs. The common initial binder had an ESMFold binder-chain mean pLDDT of 34.68. At the end of the matched search, the best BAGEL design reached 52.50, an improvement of 17.82 points, whereas the best DASTevolve design reached 74.76, an improvement of 40.08 points and a final value 22.26 points above BAGEL (Fig. 3F). The predicted complexes also differed in their realization of the intended C-terminal interface. Using a minimum heavy-atom distance threshold of 5 Å, the best BAGEL design formed four residue contacts with the ASYN C-terminal region, whereas the best DASTevolve design formed ten (Fig. 3G). The DASTevolve contact pattern was accompanied by higher confidence across much of the designed binder and a more extended predicted interface.

To distinguish program adaptation from representation, we compared Full DAST, Static DAST, Single AST, and Flat LLM from a shared initial binder over 2,000 charged candidate evaluations per arm. Apart from the defining representation or revision intervention, the arms used matched task information, prior information, LLM sampling settings, sequence-search configuration, molecular evaluation, and feasibility criteria. Static DAST retained the initial compiled program, Single AST used a unified hierarchy, and Flat LLM expressed the corresponding search guidance without the Dual-AST representation. Full DAST retained both the dual-tree representation and feedback-driven program revision. From a common initial binder pLDDT of 33.82, Full DAST reached 66.50, compared with 53.11 for Single AST, 45.54 for Static DAST, and 44.38 for Flat LLM (Fig. 3H), while best candidates passing the complete feasibility gate reached binder pLDDT values of 63.05 for Full DAST, 52.94 for Single AST, 45.54 for Static DAST, and 44.38 for Flat LLM. Full DAST also produced 204 hard-pass evaluations with binder pLDDT *≥* 50 (10.2% of the budget), compared with two for Single AST and zero for Static DAST and Flat LLM (Fig. 3J). These counts show that the higher-confidence search reached a larger set of feasible evaluations above this threshold.

Full DAST revised position-specific amino-acid rules in 18 of 19 outer updates. The records connect the search outcomes to explicit, residue-localized instructions (Table 1). Among Full DAST single-substitution children, newly targeted positions produced immediate-parent binder-pLDDT gains in 40/65 evaluations (61.5%), compared with 410/1,015 (40.4%) at other positions (Fig. 3I). Simultaneous improvement in binder pLDDT, global pLDDT, and selection loss occurred in 17/65 (26.2%) and 141/1,015 (13.9%), respectively. A generation-14 example follows the complete chain from a node-level low-confidence diagnosis through a compiled position-31 amino-acid rule to the realized M31E substitution and a 4.19-point binder-pLDDT gain. This feedback-to-action record explains how the outer loop directs local exploration and makes the tested intervention auditable. Across TrpB and ASYN, DASTevolve coupled program adaptation with effective candidate selection and improved structural-confidence outcomes. The ASYN ablations and execution traces show how the Dual-AST representation and feedback-driven revision guide sequence exploration through explicit residue policies and targeted edits, establishing a traceable connection between evaluator evidence, constrained exploration, and design outcomes.

**Table 1.** ASYN outer-loop feedback, executable policies, and sequence-level responses across controllers. Representative traces connect recorded feedback to executable policies and, for Full DAST, evaluation changes relative to the immediate parent.

| **Controller / iteration** | **Recorded diagnosis** | **Executable encoding and observed response** |
|---|---|---|
| Full DAST / 14 | “B01 pLDDT 37.33 lowest; B03 38.91 second.” | Center-node rule at residue 31 favored A/L/E/K/Q/M (weight 1.6). The executed M31E substitution increased binder pLDDT from 40.42 to 44.61 and reduced selection loss from 0.6318 to 0.6065. |
| Full DAST / 15 | “B02/B04 under-sampled per edit contract.” | A new residue-40 rule favored A/L/E/K/Q/M (weight 1.5), encoding continuity at the B04–B05 junction. H40A increased binder pLDDT from 50.41 to 52.63 and reduced selection loss from 0.6032 to 0.5925. |
| Single AST / 14 | “The lower and cterm nodes need stronger fold-stabilizing bias.” | B04/B05 policy weights were set to 1.5, with G added to their disfavored lists. Node-level amino-acid preferences changed in 13/19 updates. |
| Flat LLM / 14 | Flat sequence-search guidance | B05 used point/resampling weights 0.30/0.70 and mutation rate 0.22. Search-level amino-acid guidance changed in 19/19 updates. |
| Static DAST | Initial program retained | Compiled policies remained fixed across all 19 updates. |

### 2.3 Knowledge priors and attributed feedback guide program execution

We next examined how two complementary evidence sources affect program execution by examining two complementary forms of augmentation. In cAb-Lys3, an external VHH sequence database supplied a region-resolved empirical prior and an early feasibility gate. In APH(3^*′*^)-IIa, active-site protection and structure-based evaluators were used to distinguish a favorable scaffold-stability signal from an unfavorable ligand-pocket response. These two cases and their ablations respectively demon-strate the benefits of knowledge priors and richer evaluator feedback for protein-program optimization, complementing the architectural evaluation of DAST framework.

We selected the experimentally characterized cAb-Lys3 VHH bound to hen egg-white lysozyme (HEWL) [43]. Camelid VHHs combine a compact single-domain architecture with favorable solubility and stability, but viable designs must still respect framework and CDR-specific sequence constraints [44]. Recent sequence- and structure-generative studies have extended computational antibody engineering toward language-model-guided evolution, atomically accurate de novo design, and epitope-conditioned CDR generation [45–47]. The cAb-Lys3 system provides a reference complex, a fixed 133-residue VHH framework, a fixed 129-residue protein antigen, a protected CDR3 cysteine anchor, and paratope regions that can be evaluated independently. Therefore our search exposed 35 positions distributed across CDR1, CDR2, and CDR3 while preserving the remaining framework, the A109 cysteine, the complete HEWL sequence, and the required sequence-integrity constraints. The Structural AST restricted editing to these three CDR nodes, whereas the Functional AST separately represented core-contact recovery, broader CDR3 recovery, CDR1/CDR2 maturation, sequence integrity, and escape from the initial all-A state. In this case specification, sequence recovery denotes the fraction of predefined reference sites matching native cAb-Lys3 residue identities.

Open-ended CDR design creates many sequences that satisfy simple alphabet and length constraints but no longer resemble viable VHHs. Generic protein-language-model priors, including ProGen likelihood, provide a measure of global protein-sequence plausibility but do not explicitly capture VHH-specific regional distributions, framework–CDR compatibility, or empirical antibody constraints [12]. We therefore constructed an empirical prior from the SAbDab2 SD-H dataset, the successor to the original SAbDab resource [48, 49]. Before embedding, we removed the target structures, target antibody and antigen clusters, connected antibody–antigen components, and sequences highly similar to the target. The resulting knowledge base contained 988 unique training VHHs and 276 held-out calibration sequences. Whole-VHH, framework, CDR1, CDR2, and CDR3 sequences were encoded independently with ESM-2 `esm2_t6_8M_UR50D` [7], producing five region-specific 988 *×* 320 reference matrices (Fig. 4A). Cluster-balanced top-20 cosine similarity and held-out out-of-distribution calibration converted these embeddings into an empirical VHH plausibility score evaluated before high-cost structure prediction. This calibration is important because movement through learned sequence embeddings is not uniformly associated with biological validity [50]. CDR3 received the largest prior weight because it was the longest and least constrained design region.

**Fig. 4.**
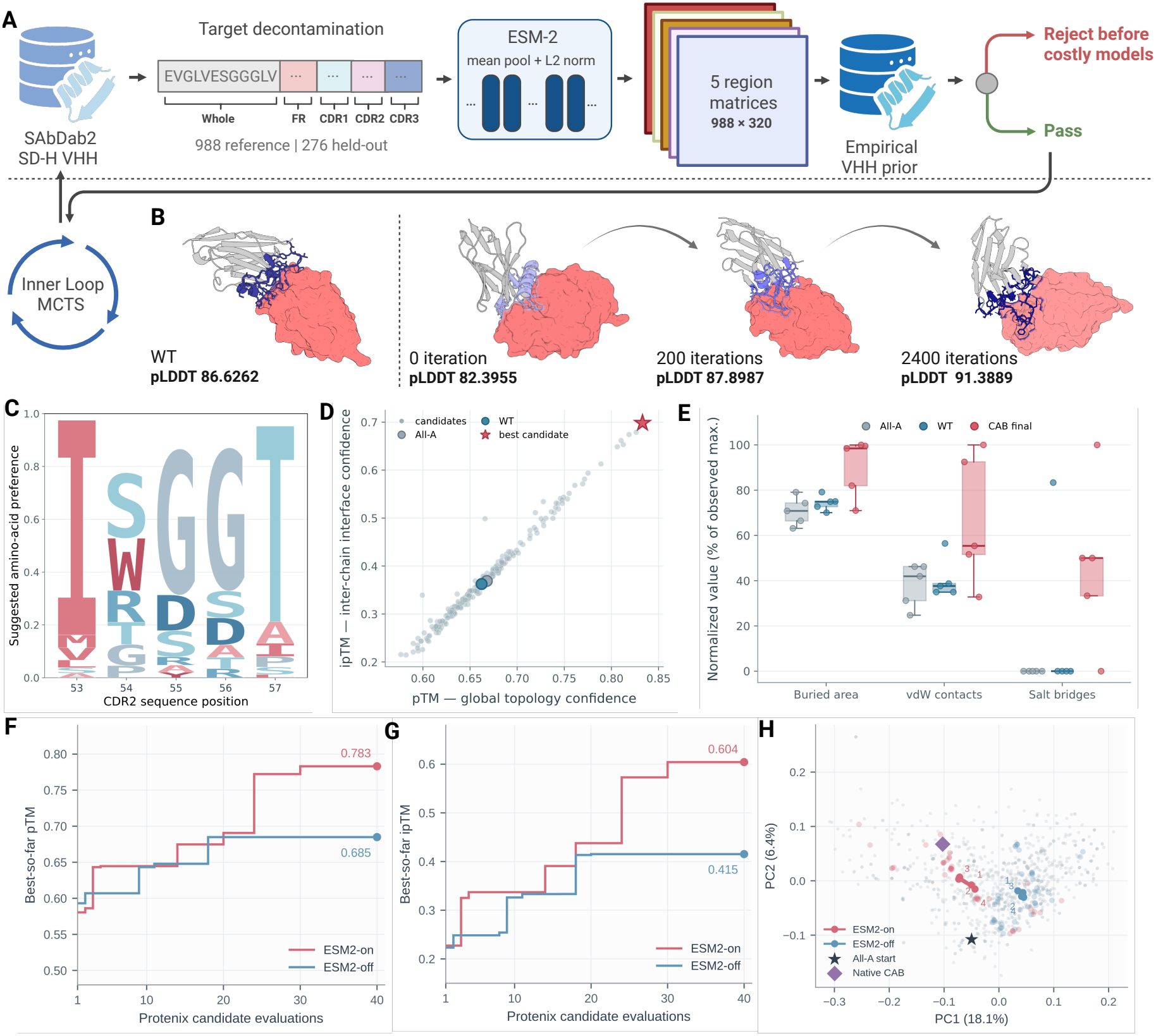
An empirical VHH prior guides cAb-Lys3–HEWL design. **(A)** Region-specific ESM-2 reference embeddings supply a calibrated VHH prior and an early plausibility gate. **(B)** Reference complex, all-A CDR starting scaffold, and designs after 200 and 2,400 iterations. **(C)** Compiled amino-acid preferences at CDR2 positions 53–57. **(D)** Candidate pTM and ipTM; the star marks the selected design. **(E)** Interface buried area, van der Waals contacts, and salt bridges across five prediction seeds, normalized to the observed maximum for each metric. Boxes show quartiles and medians; points denote seeds. **(F**,**G)** Best-so-far pTM and ipTM with ESM2-on and ESM2-off over the common initial 40-candidate window. **(H)** Archived ESM-2 sequence-space projection; numbered points mark consecutive tencandidate centroids. PC1 and PC2 explain 18.1% and 6.4% of variance. The all-A starting scaffold and native cAb-Lys3 provide reference positions.

The empirical VHH sequence prior clearly distinguished the degenerate All-A input from sequences exhibiting VHH-like characteristics. The All-A control obtained an overall calibrated plausibility score of only 0.0168 and failed the prespecified gate of 0.40, whereas our candidate reached 0.9488 and passed the gate. Structure prediction further supported the computational plausibility: the final candidate achieved the highest pTM and ipTM values among the evaluated sequences, at 0.833 and 0.698, respectively, compared with approximately 0.66 and 0.36 for the All-A controls. Across five random seeds, interface analysis yielded median values of 1,066 Å^2^ buried surface area, 103 van der Waals contacts, and three salt bridges, exceeding the corresponding values of 767 Å^2^, 78, and zero for All-A and 811 Å^2^, 70, and zero for WT, indicating that its higher model confidence was accompanied by more extensive predicted interface packing (Fig. 4E). After 2400 iterations of evolution, our best candidate achieved a pLDDT of 91.3889, outperforming the original 1JTT baseline. Notably, this improvement did not arise through convergence toward the native 1JTT sequence: our best candidate retained an overall CDR divergence of 82.9% across the three CDRs while achieving the highest pLDDT, pTM, and ipTM values. By integrating a data-driven antibody sequence prior with structure-confidence and interface-geometry evidence, DASTevolve therefore recovered a non-native, sequence-novel, and computationally credible VHH-like solution from a degenerate starting point. These results support the capacity of executable protein-program evolution to explore beyond the native sequence neighborhood and identify diverse, structurally plausible design solutions with potential for subsequent experimental evaluation.

The matched ESM2-on/off ablation comprised five rounds of 40 Protenix candidate evaluations per arm, totaling 200 evaluations for each condition. We examined both early search performance over the first 40 evaluations and the best confidence scores attained across the complete search. Both conditions used the same frozen paired and unpaired MSAs derived from a CAB WT ColabFold search. For each candidate, the query sequence was replaced while the homologous sequences and pairing assignments were retained. ESM2-on reached pTM 0.7831 and ipTM 0.6041, compared with 0.6849 and 0.4153 for ESM2-off (Fig. 4F,G). Two ESM2-on candidates jointly reached pLDDT *≥* 88, pTM *≥* 0.70, and ipTM *≥* 0.50, with the first hit at evaluation 24. ESM2-off produced no joint hit within this window. The ESM-2 projection shows distinct search directions from the shared all-A starting scaffold (Fig. 4H). The complete search trajectories also favored prior guidance: ESM2-on attained maximum pTM and ipTM values of 0.802 and 0.658, respectively, versus 0.742 and 0.518 for ESM2-off. Thus, the early advantage was accompanied by higher overall confidence endpoints. The final design combined substantial CDR divergence with high predicted complex confidence and an expanded predicted interface. Together with the matched prior comparison, this case shows how a region-specific sequence prior can be incorporated into an executable design program and guide search away from a degenerate starting sequence toward a VHH-like candidate.

We then formulated the APH(3^*′*^)-IIa design as a multi-objective enzyme-scaffold optimization task subject to functional-site preservation constraints. Using the P00552 wild-type sequence and the KAN/Mg^2+^ complex geometry from 1ND4 as references, we optimized surface and structural-support regions distal to the catalytic center while maintaining the protein length and protecting the functional core. D157, D190, N195, and D208 were designated as the Functional core: D190 acts as the catalytic base, whereas N195 and D208 participate in the reference Mg^2+^ coordination environment. We additionally fixed the KAN-contact shell and residues 44–63 associated with the ATP-donor environment. The editable sequence space was organized into five Structural-AST nodes spanning the N-lobe surface, N-lobe support, hinge support, C-lobe support, and C-terminal support regions [51, 52].

Finally, our best candidate contained 19 substitutions distributed across these five Structural-AST nodes (Fig. 5A). After globally aligning the Candidate to WT over residues 10–264, the overall back-bone RMSD was 0.551 Å. The backbone RMSDs of the Functional core and KAN-contact shell were only 0.251 and 0.359 Å, respectively, indicating that the reference functional core and KAN-binding pocket geometry were well preserved despite the introduction of distributed scaffold-support substitutions (Fig. 5B). The Protenix complex model of the Candidate was also assigned high confidence, with pLDDT, pTM, and ipTM values of 92.32, 0.951, and 0.961, respectively.

**Fig. 5.**
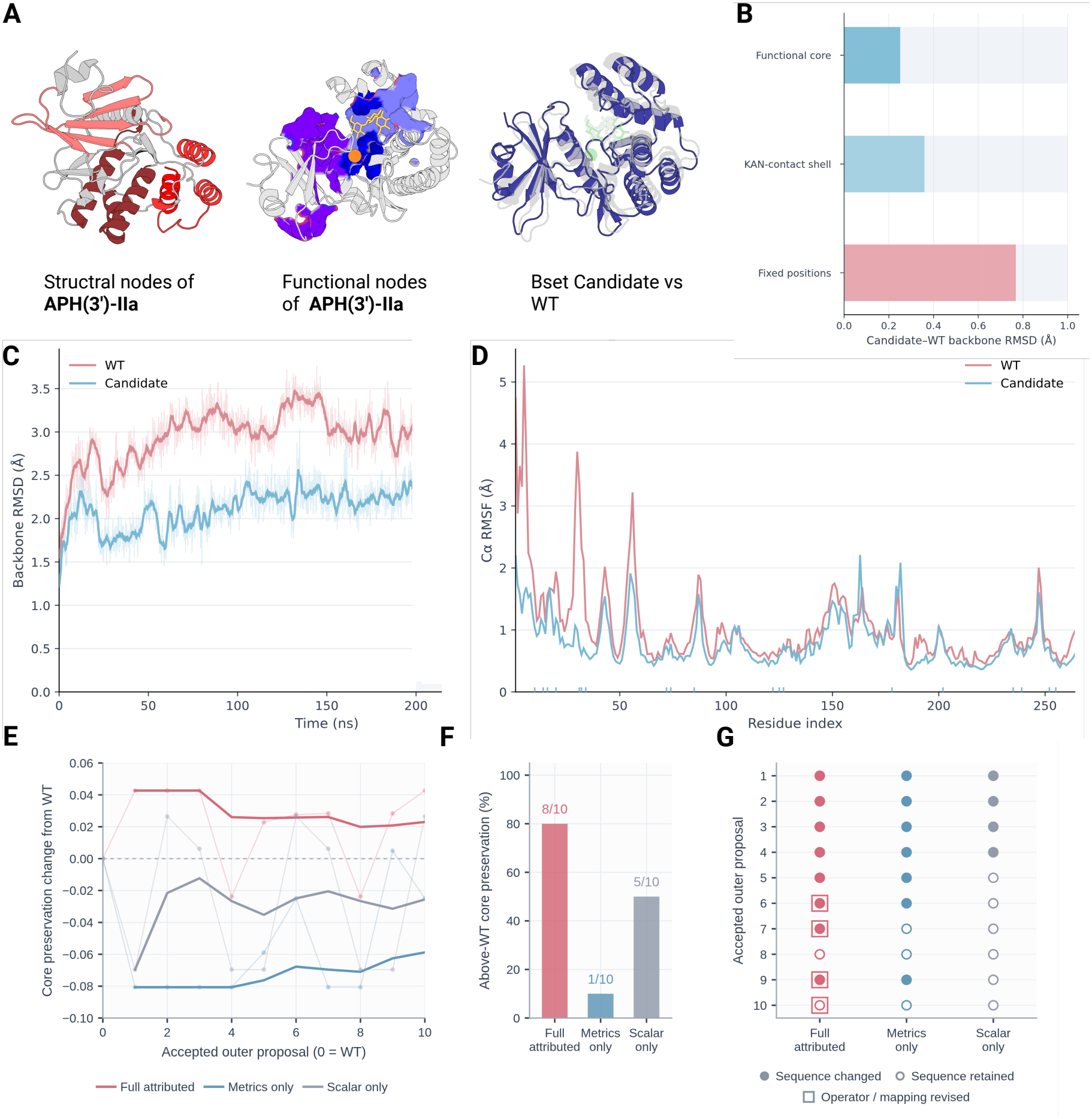
APH scaffold redesign and feedback-guided program revision. **(A)** Structural and functional nodes and the selected candidate aligned to WT. **(B)** Regional backbone RMSDs after global alignment over residues 10– 264. **(C**,**D)** Backbone RMSD relative to each starting structure (first 200 ns displayed) and per-residue C*α* RMSF over 100–300 ns. Light and dark RMSD traces show 0.1-ns samples and 21-point centered moving averages. **(E)** Corepreservation changes from each feedback arm’s own WT during ten accepted outer proposals (each outer proposal comprising 50 inner-loop iterations); thin lines show selected outcomes and thick lines show cumulative means excluding WT. **(F)** Above-WT core-preservation counts in the same window. **(G)** Parent-relative outer-loop execution: filled circles denote changed selected sequences, open circles retained sequences, and squares operator-profile or functional-mapping revisions. Proposal order follows acceptance time across the program population; each event is evaluated against its recorded parent.

To further assess dynamic stability, we performed 300-ns explicit-solvent molecular dynamics simulation for each of WT and the Candidate, using the Amber14SB protein force field, the TIP3P water model, and GAFF2/AM1-BCC parameters for KAN. Over the recorded analysis window, the Candidate exhibited a mean backbone RMSD of 2.269 Å, compared with 2.768 Å for WT, corresponding to an 18.0% reduction. The mean C*α* RMSF over 100–300 ns decreased from 1.058 Å for WT to 0.794 Å for the Candidate, a reduction of 24.9% (Fig. 5D). The mean radii of gyration were nearly identical, at 19.213 Å for WT and 19.228 Å for the Candidate over 200–300 ns, indicating that the lower structural deviation of the Candidate did not arise from substantial global collapse or expansion. Collectively, these results support a more stable and less conformationally fluctuating overall scaffold for the Candidate within the present trajectories.

We compared three feedback configurations in the APH case while retaining the shared structured-action interface and permitted operator set. Full-attributed feedback combined node- and mapping-resolved evaluator evidence with ownership and action history and cross-island context. Metrics-only and Scalar-only supplied the corresponding metric-level and scalar summaries. Each condition was represented by one retained search trajectory, and the analysis covered its first ten accepted outer proposals. The mean core-preservation change relative to each arm’s separately evaluated WT reference was +0.0230 for Full-attributed, *−*0.0588 for Metrics-only, and *−*0.0256 for Scalar-only. Eight of ten Full-attributed proposals exceeded their WT reference, compared with one and five proposals in the controls (Fig. 5E,F). These counts summarize successive observations within each search.

The retained trajectories also differed in their recorded program-level revisions (Fig. 5G; Table 2). Four Full-attributed proposals contained revisions in the analyzed program-level categories, compared with zero in either control. The selected sequence differed from its recorded parent in eight, seven, and four proposals, respectively. Scalar-only retained its parent sequence throughout proposals 5–10. Within these trajectories, the full feedback configuration combined more frequent program-level revision with higher early preservation scores. Because attribution, history, and cross-island context were supplied together, the comparison characterizes the complete feedback configurations. The archived examples additionally document how evaluator evidence was reflected in executed revisions.

**Table 2.** APH outer-loop feedback translated into executable revisions. Feedback labels abbreviate the logged evaluator fields; excerpts are verbatim fragments from the LLM’s diagnosis or hypothesis. Encodings summarize the corresponding native DesignAction fields. P denotes the accepted proposal index within each arm; outcomes are compared with that proposal’s recorded parent.

| Feedback / proposal | Feedback and LLM excerpt | Encoded revision | Recorded execution |
|---|---|---|---|
| Full attributed, P7 | Mapped fold: failed (score 0.3364).<br>“requiring the low-burden T72S substitution while rewiring site72 to global-fold measurement” | action=rewire; site72 → global-fold preservation; revert_to_wt at A34; required_mutation=S at T72. | Mapping change applied; selected A34G → T72S. Fold score 0.3364 → 0.3727; core score 0.6726 → 0.6583. |
| Metrics only, P6 | KAN pose 0.1656; fold 0.3634.<br>“Global diagnostics show weak KAN pose preservation and incomplete fold preservation” | action=revise; restore M178; required_mutation=E at D238. | Selected M178F → D238E; energy 0.3584 → 0.3099. Operator profiles and mappings retained. |
| Scalar only, P5 | final_energy=0.3585.<br>“Strong C31 parent bias combined with required retention of F178” | action=revise; retain F178; requested *p*(C31) = 0.92; required_mutation=F at 178. | Selected M178F retained; energy 0.3585 → 0.3585. Operator profiles and mappings retained. |

### 2.4 Multistate selectivity design across peptide and small-molecule recognition

Finally, we examined whether DASTevolve could coordinate selectivity or allosteric protein design across two mechanistically distinct recognition problems. The first task sought to redesign the TIAM1 PDZ domain to preferentially recognize one peptide over a closely related counter-target. The second sought to reshape the ligand-recognition and allosteric-coupling landscape of the transcriptional regulator TtgR toward naltrexone while retaining the apo fold and dimeric architecture. These cases examine how a shared executable representation coordinates state-specific objectives with structural constraints across peptide and small-molecule recognition.

PDZ domains typically recognize short C-terminal peptide motifs to form family-specific interaction networks [53]. In its native state, the TIAM1 PDZ domain exhibits limited discrimination between the Caspr4 peptide, ENQKEYFF-COO^*−*^, and the Syndecan-1 peptide, TKQEEFYA-COO^*−*^. Experimentally, WT binds Caspr4 and Syndecan-1 with dissociation constants of 19.0 and 26.9 *µ*M, respectively, corresponding to a selectivity ratio of only 1.42. A previously reported quadruple mutant increased this ratio to 6.67, primarily by weakening Syndecan-1 binding [54]. These experimental measurements define the design context, and the computational comparisons below concern the newly designed candidate in its two peptide-bound states. We therefore formulated TIAM1 redesign as an explicit positive–negative task in which the Caspr4-bound state was favored, the Syndecan-1-bound state was penalized, and both the apo fold and the terminal-carboxylate recognition geometry shared by the two peptide states were preserved.

The 90-residue TIAM1 construct was decomposed into 17 non-overlapping structural nodes and 10 overlapping functional nodes (Fig. 6A). At each outer-loop revision, the executable program specified editable structural regions and state-specific guidance for inner sequence search in the apo, Caspr4-bound, and Syndecan-1-bound states. The active mutation set was restricted to four to eight positions per round. This representation allowed mutations to be introduced around the peptide-recognition surface while protecting the fold-defining core and the conserved interaction geometry shared by both peptide states. Eventually, the highest-ranked feasible program produced the candidate with a final normalized objective value of 0.1621. Its predicted apo structure retained high confidence, with a pLDDT of 85.75 and a pTM score of 0.901. In the peptide-bound states, the predicted interface TM score was 0.941 for Caspr4 and 0.765 for Syndecan-1, yielding an ipTM difference of 0.176 between the candidate’s Caspr4-bound and Syndecan-1-bound states. Consistently, the mean symmetric peptide– protein cross-interface PAE was far lower for Caspr4 than for Syndecan-1 (3.10 versus 6.17 Å; Fig. 6B). We further performed 300-ns explicit-solvent simulations of the final candidate in complex with either Caspr4 or Syndecan-1. Over the analysis interval, the mean PDZ C*α* RMSD was lower in the Caspr4 complex than in the Syndecan-1 complex (0.2001 versus 0.2717 nm), corresponding to a reduction of 26.35%. During the final 100 ns, this difference increased to 34.57% (Fig. 6C). Interface and peptide-geometry measurements provided complementary evidence (Fig. 6D). Relative to the Syndecan-1 complex, the Caspr4 complex exhibited 3.11% more interfacial atom contacts and 19.84% more contacting PDZ residues per frame over 300 ns, indicating broader coverage of the protein– peptide interface. The mean PDZ–peptide center-of-mass distance was 5.77% shorter, and the peptide radius of gyration was 11.42% lower over the complete 300-ns trajectory, consistent with a peptide position closer to the PDZ domain and a more compact bound conformation. The Caspr4 advantage in the number of contacting PDZ residues remained approximately 17.7–19.8% across the tested contact-distance cutoffs, supporting the robustness of this trend to the specific cutoff choice. Together, the predictions and sampled trajectories showed distinct structural signatures for the candidate in its two peptide-bound states. The Caspr4-bound state combined higher predicted interface confidence, lower PDZ backbone deviation, broader interface-contact coverage, shorter peptide–domain separation, and a more compact peptide conformation. These state-specific signatures are consistent with the intended computational design objective.

**Fig. 6.**
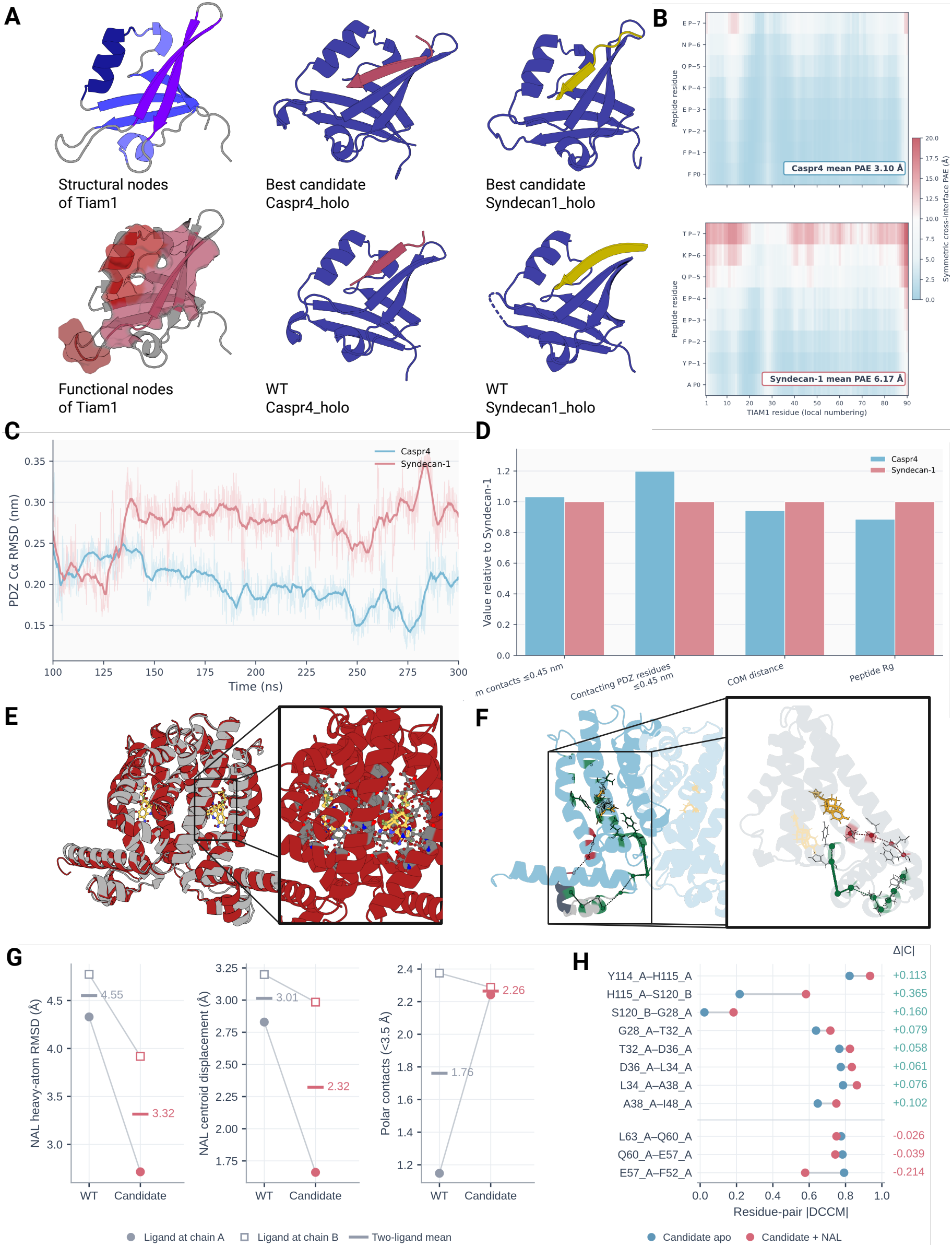
Multistate design of TIAM1 PDZ and TtgR. **(A)** TIAM1 structural/functional nodes and candidate/reference peptide complexes. **(B)** Symmetric cross-interface PAE for Caspr4 and Syndecan-1. **(C)** Candidate PDZ C*α* RMSD in the two peptide-bound states; light curves show 0.1-ns samples and dark curves 5.1-ns moving averages. **(D)** Interface and structural metrics normalized to Syndecan-1. **(E**,**F)** TtgR pocket redesign and selected residue-correlation routes. Green and red in F indicate stronger and weaker absolute correlations in candidate/NAL relative to candidate apo. **(G)** WT/NAL and candidate/NAL ligand-localization metrics. Circles and squares denote chain-A and chain-B ligand means; short horizontal marks indicate their average. RMSD and centroid displacement use each trajectory’s starting ligand coordinates after associated-chain alignment. Polar contacts count N/O pairs within 3.5 Å. **(H)** Absolute C*α* DCCM correlations for all 11 selected correlation edges in candidate apo and candidate/NAL trajectories; Δ|*C*| = |*C*_NAL_| *−* |*C*_apo_|.

We lastly considered a different form of multistate selectivity: redesigning the TetR-family transcriptional regulator TtgR toward naltrexone recognition. Structural studies have shown that TtgR contains distinct, partially overlapping ligand-binding sites that support chemically diverse effectors [55]. More recently, high-throughput screening and a naltrexone-bound variant structure demonstrated that this malleable pocket can support non-native ligand responses [56]. The Functional AST treated naltrexone as the target ligand, included endoxifen and tamoxifen as relative counter-states, and retained separate constraints for the apo fold, homodimer organization, and DNA-binding domain. The 210-residue monomer was represented by 17 structural nodes. The editable region covered the ligand pocket and the intervening helices connecting ligand recognition to the N-terminal DNA-binding module.

The final candidate contained 12 substitutions distributed across three spatially connected modules. The TtgR molecular-dynamics analysis comprised three systems: WT/NAL, candidate apo, and candidate/NAL, with two NAL molecules in each ligand-bound system and one trajectory per system. Over 20–220 ns, naltrexone heavy-atom RMSD decreased from 4.551 Å in the wild-type complex to 3.316 Å in the candidate, and ligand–pocket center-of-mass displacement decreased from 3.014 to 2.323 Å (Fig. 6G; Table 3). The number of polar distance contacts increased from 1.761 to 2.265, whereas retention of contacts present in the initial structure changed only slightly. These changes correspond to 27.1% lower ligand RMSD, 22.9% lower centroid displacement, and 28.6% more polar contacts, averaged across the two ligand sites over 20–220 ns. The contact increase was driven mainly by chain A, with a small decrease at the chain-B site. Both displacement measures use each trajectory’s starting ligand position after alignment of the associated protein chain. These results show reduced NAL displacement from the respective starting poses and an altered polar-contact environment in the candidate trajectory relative to WT/NAL.

**Table 3.** Ligand-localization and regional motion-correlation metrics for TtgR. Columns compare WT/NAL with candidate/NAL, and percentage changes use WT/NAL as the reference. Figure 6F,H separately compares selected residue-pair correlations between candidate apo and candidate/NAL.

| **Metric** | **WT/NAL** | **Candidate/<br>NAL** | **Δ (%)** |
|---|---|---|---|
| **Ligand localization and pocket contacts** |  |  |  |
| NAL heavy-atom RMSD (Å) | 4.551 | 3.316 | −27.1 |
| NAL–pocket COM displacement (Å) | 3.014 | 2.323 | −22.9 |
| Polar distance contacts (< 3.5 Å) | 1.761 | 2.265 | +28.6 |
| Initial-contact retention | 0.634 | 0.642 | +1.3 |
| **Regional motion correlations** |  |  |  |
| Node-to-rest mean \|DCCM\| | 0.1487 | 0.1641 | +10.4 |
| Cross-chain mean \|DCCM\| | 0.1183 | 0.1495 | +26.4 |

NAL, naltrexone; COM, center of mass; DCCM, dynamic cross-correlation matrix. Ligand-localization and contact metrics are averaged over the two ligand sites within each trajectory.

The candidate apo and NAL-bound trajectories differed in their conformational dynamics. Candidate dimer C*α* RMSD was 3.320 Å in the apo trajectory and 2.638 Å in the NAL-bound trajectory, while the RMSD of residues 70–120 was 2.319 and 2.004 Å, respectively. The RMSF of this region was higher in the NAL-bound trajectory, increasing from 0.878 to 1.066 Å. Thus, the NAL-bound trajectory combined lower global and regional backbone deviation with greater local fluctuations in this region.

Comparison of the candidate apo and NAL-bound trajectories revealed differences in selected residue-motion correlations (Fig. 6F,H). Mean absolute DCCM correlation across the eight edges of the selected route spanning the two subunits increased from 0.5842 to 0.7111, whereas the corresponding mean across the three edges of the selected intrachain route decreased from 0.7837 to 0.6906. Eight of the 11 selected edges showed higher absolute correlation in the NAL-bound trajectory, and three showed lower values. The largest increase occurred between H115_A_ and S120_B_ (Δ|*C*_*ij*_| = +0.3651), whereas the largest decrease occurred between E57_A_ and F52_A_ (Δ|*C*_*ij*_| = *−*0.2143). These differences motivate a residue-resolved allosteric hypothesis involving altered coordination between the ligand-proximal and DNA-binding regions, with specific residue pairs nominated for subsequent perturbation and ligand-dependent DNA-binding assays.

Together, these cases demonstrate how executable protein programs coordinate multistate design objectives through explicit state definitions, structural constraints, and state-specific evaluation. The TIAM1 candidate exhibited differential interface confidence and distinct contact and conformational profiles in its two peptide-bound states. The TtgR candidate showed reduced NAL drift relative to WT/NAL and differences in selected residue-motion correlations between its apo and NAL-bound trajectories. These results connect multistate program execution to concrete candidates, traceable computational evidence, and residue-resolved hypotheses for testing selectivity and allosteric regulation.

## 3 Discussion

Sequence generators, structure predictors, and molecular evaluators provide the computational foundation of protein design. Applying these models to functional design under multiple constraints requires explicit requirements, defined intervention boundaries, and search strategies that can be revised in response to evidence. DASTevolve addresses this process-level problem by extending iterative optimization from candidate molecules to the design program directing molecular search, providing an executable basis for program-level self-iteration in targeted functional design. A large language model reasons over functional requirements, structural regions, and intervention hypotheses, while specialized molecular models propose and evaluate candidates. Molecular feedback then informs revisions to the program governing subsequent search. Design hypotheses thus acquire an explicit, machine-readable representation and can be assessed through concrete operations and their molecular consequences. This organization brings the intervention strategy itself within the scope of optimization, allowing design programs and candidate molecules to participate jointly in iterative exploration.

Our study provides a common representational and executable framework for constrained optimization under finite evaluation budgets, adaptation to changing objectives, coordination of global properties with local functional requirements, and design involving competing states and preservation requirements. In the matched-budget binder-design comparison, DASTevolve improved predicted structural confidence and target-region contacts and ablation experiments further supported contributions from structured representation and feedback-driven revision. Objective-switch and feedback experiments showed that reference information and attribution-rich evaluation evidence can influence program-level decisions and subsequent search outcomes. The multistate tasks illustrated how a single representation can organize mutually constraining requirements and translate them into interventions and evaluation rules. Together, these findings support explicit task representation and evidence-driven program revision as a route to task-level adaptation. Their application across the examined tasks also supports reuse of the framework and its representation across design scenarios. This mechanism builds on organizing hierarchical design requirements as syntax trees and compiling node-level constraints into molecular search objectives [33], together with principles of composable objectives, model composition, and tool orchestration [28, 34, 35]. DASTevolve extends these principles by explicitly linking functional requirements to structural permissions through the Dual-AST and making these associations revisable within prescribed boundaries. The functional AST organizes properties to promote, suppress, or preserve, while the structural AST defines intervention regions and permissible operations. Their typed mappings provide an interface for associating evaluation evidence with specific design hypotheses. This separation allows a functional requirement to involve multiple structural regions and a region to participate in several requirements, preserving these relationships during constrained program revision. From a programming-language perspective, the Dual-AST is an executable intermediate representation linking biological hypotheses to molecular search: the outer loop revises hypotheses and execution configurations, compilation resolves them into operational scopes and constraints, and the inner loop searches for candidates within those constraints. This organization is relevant when improvements in aggregate metrics coexist with adverse responses in regions associated with functional preservation. When global metrics and local responses conflict, this representation helps identify the implicated requirements and regions and informs adjustments to intervention strategies. Within each task phase, shared outer-loop evaluation criteria allow competing programs to be compared on a consistent scale. The Dual-AST therefore gives the hypothesis–operation–evidence–revision cycle explicit semantics and constraints, making program-level self-evolution inspectable and traceable. Current evidence supports problem detection and attribution more strongly than reliable automatic correction.

Obviously, current designed candidates lack prospective experimental validation, program revision depends on molecular-evaluator reliability, and the present evidence establishes task-level adaptation without demonstrating general, transferable self-improvement. Evaluator biases may misdirect subsequent revisions, while sensitivity to model version, context, and sampling configuration requires further assessment. Therefore, the decisive next test is whether program-guided revision improves experimental outcomes in prospective design–build–test–learn cycles [57, 58]. Proposed extensions address feedback exchange, reusable search guidance, and uncertainty handling, and the associated neural search-guidance scheme is formalized in Supplementary Information Section S1. The conceptual contribution is to bring candidate molecules and their design logic into a common process of testing and revision. By linking functional intent, intervention strategies, and molecular consequences, this framework makes the design program itself a scientific object that can be evaluated, revised, and reused.

## 4 Methods

The framework overview of DASTevolve is presented with Figs. 1 and 2 in Results. This section specifies its executable representation, program-revision rules, inner and outer optimization procedures, and backend interfaces.

### 4.1 Executable DAST representation

DASTevolve represents a protein-engineering task as an executable and evolvable protein program

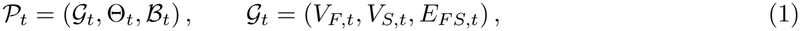

where *t* denotes the outer-loop generation. The Dual Abstract Syntax Tree (Dual-AST) *G* _*t*_ contains functional nodes *V*_*F,t*_, structural nodes *V*_*S,t*_, and typed functional–structural mapping edges *E*_*FS,t*_. The search configuration Θ_*t*_ specifies the inner optimization policy, whereas the edit envelope *ℬ*_*t*_ defines which aspects of the current program may be revised in the next outer iteration.

A functional node specifies a biological state, an objective or hard constraint, the desired optimization direction, an evaluator binding, and a policy for missing evidence. Its state is classified as positive, negative, or preserve, corresponding respectively to an outcome that should be promoted, suppressed, or maintained. Functional nodes therefore describe *what* should change or remain invariant without directly prescribing an amino-acid substitution.

A structural node specifies a chain-specific residue selector, whether the selected region is editable or frozen, the invariants associated with that region, and the sequence-intervention actions that are permitted. Editable nodes expose one or more actions with explicit mutation budgets, whereas frozen nodes admit no sequence-changing action and preserve the parent sequence.

Each mapping edge connects a functional node *f*_*i*_ to a structural node *s*_*j*_ through an allowed action *a*_*ij*_:

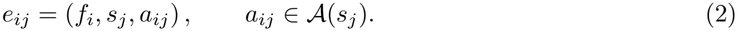

The edge has the semantic relation *realizes*: it states that applying action *a*_*ij*_ within the residue scope of *s*_*j*_ constitutes a testable intervention for functional objective *f*_*i*_ (Fig. 2A). This relation provides a lineage

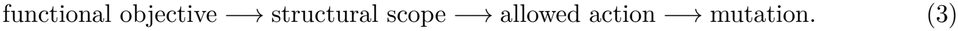

The executable schema requires unique node and edge identifiers, valid residue bounds, no dangling mappings, no mapping into a frozen node, compatibility between each edge operator and its target structural node, and at least one mapping for every active functional node. Programs that violate these requirements are not executable.

### 4.2 Protein program revision and compilation

The outer loop operates at the program level and refines the design specification that guides sequence optimization. Given a parent program *P*_*t*_, proposal *k* produces a typed revision *δ*_*t,k*_. The revision is accepted only if it lies within the edit space permitted by the parent envelope and passes schema, identity, permission, residue-bound, invariant, and evaluator-binding checks:

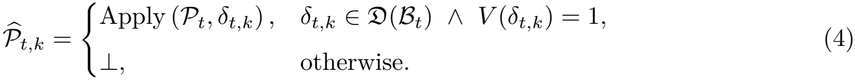

*D*(*ℬ*_*t*_) is the set of program modifications authorized by the parent envelope. Depending on the case, an authorized revision may resize or relocate an editable selector, change an action profile or mutation budget, modify a functional–structural mapping, strengthen negative-design emphasis, or adjust the inner-search policy. The case-owned outer-evaluation specification, protected residues, immutable reference sequences, and mandatory feasibility gates are outside the program-revision edit space. An invalid proposal returns ⊥ and does not initiate an inner search. A valid revised program is compiled into an executable search contract

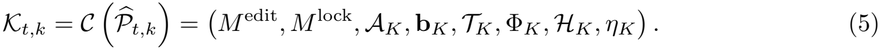

Here, *M* ^edit^ and *M* ^lock^ are the concrete editable and protected residue masks, *A*_*K*_ is the set of permitted sequence interventions, **b**_*K*_ contains mutation budgets and position policies, *T*_*K*_ contains state-specific sequence or structure templates, Φ_*K*_ binds evaluator terms to functional nodes, *ℋ*_*K*_ contains the effective hard gates, and *η*_*K*_ is a content-derived contract identity.

For a sequence bundle **x** and immutable reference **x**^ref^ , define the mutation set

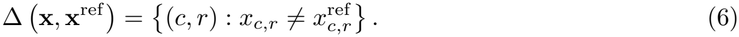

The compiled contract induces the sequence space

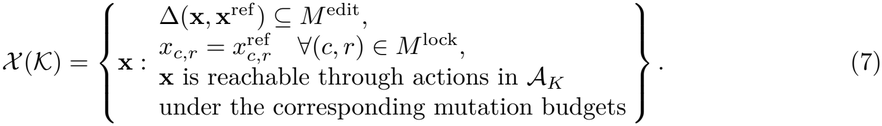

Compilation determines the inner-search scoring configuration, reachable sequences, protected residues, and permitted interventions, while retaining the case-owned outer-evaluation specification and mandatory biological requirements. An outer-loop revision can consequently change the geometry of the inner sequence-search space even when the reference sequence remains unchanged (Fig. 2B).

### 4.3 Outer loop: Protein Program Evolver

We distinguish the program-dependent objective used to guide inner sequence search from the case-owned criterion used to compare competing programs. Let Ξ denote the shared outer-evaluation specification, including the assessed molecular states, evaluator definitions, score normalization, aggregation weights, evidence-availability policies, and mandatory feasibility gates. Within a given task and evaluation phase, this specification is fixed across outer-loop iterations and is not part of the AST edit space.

For a fixed valid program *P*, the inner optimizer approximately solves

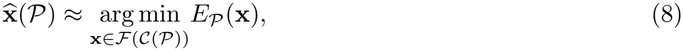

where *ℱ* (*C* (*P*)) is the gate-passing subset of the contract-valid sequence space and *E*_*P*_ is the inner-search loss associated with the compiled program. Each returned candidate is assessed using the common outer criterion *E*_out_(·;Ξ). In an outer-aligned configuration, the two criteria can coincide and the same evaluation evidence can be reused. The outer problem is therefore

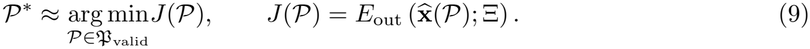

Thus, a program affects its outer score through the candidate produced by its constrained inner search, rather than by redefining the criterion used to judge that candidate. Outer performance ranking uses the shared case-owned feasibility requirements and evaluation specification, and all-infeasible runs remain explicitly marked as such. This constrained bilevel formulation separates program evolution from sequence optimization while keeping cross-program evaluation comparable (Fig. 2C). Both levels operate over discrete spaces under finite computational budgets.

For every proposal, the outer-loop optimizer receives the parent program, evaluator residuals, node-level evidence, failure diagnoses, and the current edit envelope. The resulting candidate revision is submitted to the validator. Selection then operates on its effective compiled phenotype, as defined by the resolved residue masks, permitted actions, evaluator configuration, and search policy.

The population layer follows the evaluator-guided program-evolution paradigm exemplified by AlphaEvolve [36] and adapts OpenEvolve’s open-source population and island-management mechanisms [39] to typed protein programs. Let 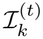 denote the population maintained by island *k* at outer iteration *t*, and let *B*_*k*_ denote its population capacity. All islands share the same program schema, case-owned feasibility requirements, and outer objective vector. The components of **J**(*P*) are evaluated under the same specification Ξ; *J* (*P*) in Eq. 9 denotes the corresponding case-defined scalar aggregate. Each island applies a configurable role projection

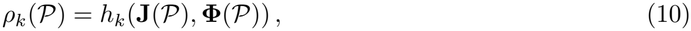

where *h*_*k*_ emphasizes a task-specific search responsibility, such as interface and fold quality, positive–negative state separation, exploration of alternative structural regions, or robustness against unfavorable states. The projection *ρ*_*k*_ supplies a secondary selection pressure among candidates with comparable feasibility and outer-level performance. It does not alter the shared validation rules, measurement definitions, normalization, or case-owned objectives. Each role projection is applied consistently to all programs considered by that island, and migrated programs are assessed under the destination island’s selection rule.

If 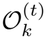 is the set of candidate programs generated for island *k*, its population update can be written as

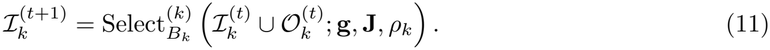

Here, 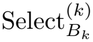 first enforces program validity and case-owned feasibility, then considers multi-objective performance and the role-specific projection when retaining at most *B*_*k*_ programs. Consequently, island specialization affects parent and survivor selection while preserving a common biological specification.

At migration iterations *t ∈ τ* ℕ, an eligible subset 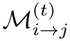 is selected from a source island *i* and submitted to a complementary target island *j*. Admission isdetermined by the target island’s selection operator:

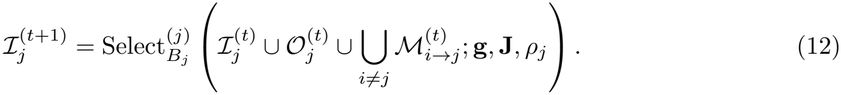

Migration therefore constitutes target-specific reconsideration. The transferred record contains the versioned program, its compiled phenotype, evaluation results, and provenance. Program-specific MCTS trees remain local to their compiled contracts: each accepted program initializes a fresh inner tree, and visit counts and value statistics are not shared across different contracts. This separation allows islands to exchange useful semantic revisions while maintaining distinct optimization trajectories and search responsibilities.

### 4.4 Inner loop: Protein Sequence Optimizer

For a fixed search contract *K*, each MCTS node *v* contains a concrete sequence bundle **x**_*v*_ *∈ X* (*K*). A tree edge represents a typed intervention *a* drawn from the actions permitted by the current contract:

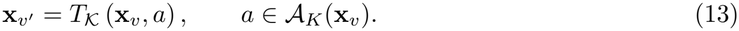

A child is materialized only after sequence-validity, mutation-budget, ownership, and invariant checks. Duplicate sequence proposals are short-circuited before expensive provider evaluation and do not consume expansion capacity or trigger additional backpropagation.

Because the number of possible mutation actions can be large, progressive widening [59] limits the number of committed children for parent *p* and expansion context *k*:

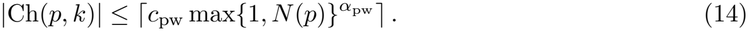

For 0 *< α*_pw_ *<* 1, the exposed branching factor grows sublinearly with the number of visits. The search can therefore revisit promising branches while gradually exposing new structural regions, operators, or proposal contexts.

Among the exposed children, tree selection uses a prior-weighted upper-confidence rule in the PUCT family [60, 61]:

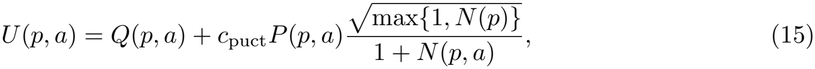

where *Q*(*p, a*) is the current mean reward, *P* (*p, a*) is the proposal prior, *N* (*p*) and *N* (*p, a*) are visit counts, and *c*_puct_ controls exploration. Protein-language-model and node-level information can contribute to the proposal prior, but these priors are not treated as direct measurements of task fitness.

Let *L*_0_ denote the root selection loss and 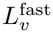 the loss assigned to candidate *v* after the configured fast evaluation. The bounded search reward is

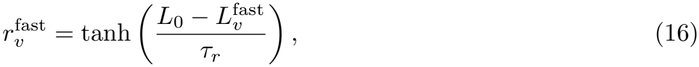

where *τ*_*r*_ *>* 0 is the reward scale. For every node *u* on the path from the root to *v*, ordinary backpropagation updates

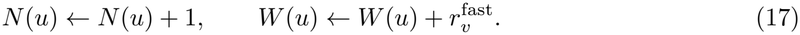

At configured checkpoints, selected candidates receive a higher-fidelity structural evaluation. If the upgraded reward for node *v* differs from its currently assigned reward, the correction is

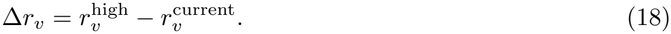

This correction is propagated along the existing ancestral path:

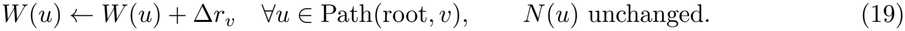

Higher-fidelity evidence updates the value estimates of the active tree while leaving its visit counts unchanged. These updated estimates inform subsequent tree selection, allowing the high-fidelity oracle to provide online value correction throughout the search (Fig. 2D).

### 4.5 Evaluator reports and memory updates

Evaluator outputs are divided into soft optimization terms, hard feasibility gates, and diagnostic measurements. Let *s*_*i*_(**x**) *∈* [0, 1] be a normalized score with non-negative weight *w*_*i*_. Define an inclusion variable *z*_*i*_ and effective residual 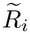 as

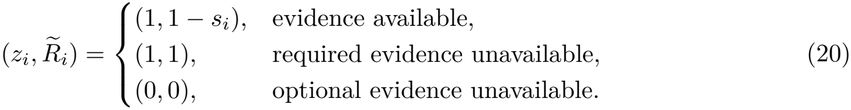

The evaluator energy is

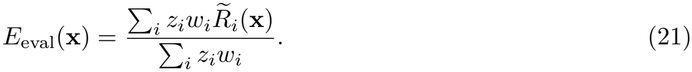

Required missing evidence therefore receives the worst residual, whereas explicitly optional missing evidence is excluded. The enabled terms can include sequence plausibility, structural confidence, interface quality, clash burden, multistate objectives, and task-specific measurements. Their identities and weights are case dependent. For cross-program comparison, these terms, their normalization, their weights, and their missing-evidence rules are supplied by the common specification Ξ, yielding *E*_out_(**x**; Ξ). Program-specific choices used to guide inner search do not redefine this outer criterion.

Hard gates are evaluated separately. Let *g*_*ℓ*_(**x**) *∈*{0, 1} denote the decision of gate *ℓ*. The feasible survivor set is

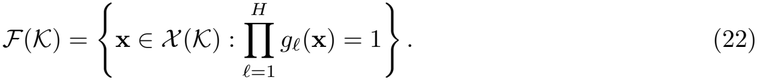

When this set is non-empty, the candidate returned by the inner search is

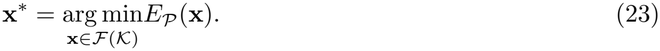

Thus, hard feasibility is applied before objective ranking. A severe violation cannot be compensated by improvement in an unrelated term. If all candidates are infeasible, the implementation returns a diagnostic fallback based on hard-gate coverage and margins, but explicitly marks the result as all-infeasible.

The evaluator report records raw measurements, normalized scores, evidence availability, feasibility outcomes, residual decomposition, node attribution, and failure diagnoses. Reporting and persistent memory have different scopes: reports can describe failed or incomplete candidates, whereas memory can receive only eligible evidence associated with the selected winner. Inner-tree visits and transient rewards remain local to the current contract execution.

Finally, content hashes and generation records preserve the causal chain

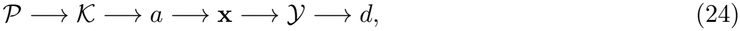

where *Y* denotes the evaluator evidence and *d* denotes the feasibility and selection decision. This lineage connects each reported sequence to the program, contract, and intervention under which it was generated.

### 4.6 Model backends

The executable contract is model-agnostic and accesses molecular backends through standardized interfaces that define their input schemas, output fields, availability checks, and failure states. Users can select, replace, or extend these backends through case-level configuration and adapter registration, provided that each implementation conforms to the corresponding interface schema and output semantics.The sequence-scoring interface provides model-derived evidence for candidate generation and ranking. Autoregressive protein language models such as ProGen and ProGen2 supply sequence log-likelihoods, whereas ESM-2 supplies masked-residue probabilities, sequence representations, and region-resolved embedding priors [7, 12, 62]. These outputs enter the search as sequence-plausibility scores, mutation-proposal priors, or representation-level evidence, with their model identity and provenance retained throughout optimization.

The structure-prediction interface maps candidate sequences and molecular-state specifications to standardized structural artifacts and confidence annotations. ESMFold and ESMFold2 provide comparatively low-cost single-chain and complex-capable predictions [7, 63], while AlphaFold 3 and Protenix-family backends, including Protenix-v2, support higher-fidelity multichain and multicomponent structure evaluation [5, 11, 64]. The case configuration assigns the required backend and fidelity tier to each evaluator call. Predicted coordinates, confidence measures, backend versions, and availability states are then returned through a common result schema, allowing structure predictors to be exchanged without changing the protein-program representation or compiled contract semantics. A separate evaluator interface applies case-configured structural, energetic, interaction, and dynamical analyses to the resulting artifacts. The PAE-derived ipSAE score supplies interface-confidence evidence [65], whereas DockQ measures the quality of a predicted complex when an experimental or designated reference structure is available [66]. PyRosetta supports scriptable Rosetta-based structural scoring and local modeling operations [67], and FoldX provides rapid empirical estimates of complex interaction energies [68]. GetContacts extracts residue-level non-covalent interactions and trajectory contact occupancies [69], which can be combined with geometric and dynamical analyses. Each output is retained as a distinct evidence channel with its native definition, directionality, fidelity, availability state, and provenance in the evaluator report. Additional evaluation tools can be registered through the same adapter mechanism, allowing users to configure the evidence channels required by each engineering case while preserving standardized reporting and provenance.

## Data Availability

The source code for DASTevolve, including the Dual-AST representation, program-compilation and evolution workflow, MCTS-based search, evaluator interfaces, and associated configuration files, is available on GitHub at https://github.com/YiyangZHA0/DASTevolve. LLM providers, API endpoints, model versions, sampling parameters, and molecular-evaluation backends can be configured or replaced through modular interfaces.

Protein sequences, canonical residue numbering, and functional annotations used to initialize the case specifications were obtained from UniProt (https://www.uniprot.org/). Experimentally determined protein structures, protein–protein complexes, protein–peptide complexes, and ligand-bound structural templates were obtained from the RCSB Protein Data Bank (https://www.rcsb.org/). The corresponding UniProt accession codes and PDB identifiers are recorded in the case-specific configuration and evidence files. The TrpB benchmark was constructed from the experimentally measured combinatorial sequence–fitness landscape reported by Johnston et al. [40]. The original landscape and associated data are publicly available through the authors’ data record at https://doi.org/10.22002/h5rah-5z170. The external antibody-sequence knowledge base used for the cAb-Lys3 case was derived from the June 2026 SAbDab2 SD-H release, the successor to the original SAbDab resource [48, 49].

Third-party software, pretrained model weights, experimental datasets, and external databases remain subject to their original licenses, access conditions, and redistribution policies.

## Acknowledgements

This work was financially supported by the National Key Research and Development Program of China (2024YFA1306400), the Zhejiang Province Pioneer Plan (2025C01189), and the Huadong Medicine Joint Fund of Zhejiang Provincial Natural Science Foundation of China under Grant No. LHDMZ25H300001.

